# Label-free Pathogen Identification with Microscopy Imaging and Deep Learning

**DOI:** 10.64898/2026.06.22.733661

**Authors:** Xudong Zhang, Tongtong Zhou, Siqi Guo, Wenbin Du, Zhou Tong, Jiajia Zheng, Ning Shen, Junhao Zhu, Jun Wang

**Affiliations:** Laboratory of Pathogen Microbiology and Immunology, Institute of Microbiology, Chinese Academy of Sciences, Beijing, China; University of Chinese Academy of Sciences, Beijing, China; Department of Laboratory Medicine, Peking University Third Hospital, Beijing, China; Department of Pulmonary and Critical Care Medicine, Peking University Third Hospital, Beijing, China

**Author notes:** Corresponding to: JW JZ. These authors contributed equally (Xudong Zhang, Tongtong Zhou).

## Abstract

Rapid and accurate pathogen identification is crucial for the clinical management of infectious diseases, particularly sepsis and severe respiratory infections, yet standard clinical workflows remain slow and resource-intensive. Here, we developed an automated, high-throughput imaging platform built on standard, clinically accessible bright-field microscopy, and generated a large dataset comprising 24.9 million label-free bacterial cells across six focal pathogens. Leveraging this resource, we trained a neural network (ESKAPe-ResNet) to identify ESKAPe species at the single-bacterium level. The model achieved >92% accuracy in species-level classification and >82% accuracy in quantifying ESKAPe abundance in mock mixtures, with high specificity against non-ESKAPe bacteria. In clinical validation using sputum, bronchoalveolar lavage fluid and blood samples from patients with respiratory infections and sepsis, the approach correctly identified the dominant ESKAPe pathogen in >78% of samples after minimum broth culture enrichment. The imaging-to-identification pipeline was completed in under 10 minutes, and coupled with brief cultivation, the median time to accurate identification was reduced to 5–6 hours, compared with days for conventional blood culture-based workflows. This work establishes the proof-of-principle for label-free, hardware-minimal rapid pathogen identification, providing a clinically deployable workflow to expedite diagnosis and reduce mortality in severe bacterial infections.

## Introduction

Sepsis and severe respiratory infections caused by bacterial pathogens, particularly the multi-drug resistant ESKAPE group—*Enterococcus faecium*, *Staphylococcus aureus*, *Klebsiella pneumoniae*, *Acinetobacter baumannii*, *Pseudomonas aeruginosa* and *Enterobacter spp*.—pose major global health threats and are associated with substantial mortality in intensive care settings^1–3^. These organisms encompass multiple critical priorities designated by the World Health Organization (WHO) in its Priority Pathogens List for research and development of new antibiotics ^4,5^. Rapid and accurate identification of the causative pathogen is crucial for guiding therapy in severe infections, especially in patients with respiratory infections or sepsis in the intensive care unit (ICU); it has been estimated that each one-hour reduction in time to appropriate antimicrobial therapy is associated with a 7% decrease in mortality^6^.

Currently, the standard clinical workflow of bacterial pathogen identification typically requires 12-72 hours of culture incubation^7^, followed by species identification, most commonly using matrix-assisted laser desorption ionization-time of flight mass spectrometry (MALDI-TOF MS) where available^8^. While being extensively validated in the clinic, this workflow remains prone to false-negative results and is both resource- and labor-intensive. Recent advances in metagenomic or targeted next-generation sequencing and nanopore-based approaches have considerably shortened time to result^9,10^; nevertheless, besides the requirement of sequencer and accompanying chemistry, these workflows typically require 6–24 hours and remain susceptible to false-positives introduced by background contaminations, low biomass sampling, and host DNA depletion inefficiencies^11^. Thus, a diagnostic approach that simultaneously delivers speed, accuracy, and broad clinical accessibility remains an unmet need^12^.

This unmet need has renewed interest in microscopy-based strategies^13^. Light microscopy has been a cornerstone of clinical microbiology for over a century and remains ubiquitous across clinical laboratories, yet its diagnostic resolution has long been limited by the human eye’s poor ability to interpret subtle morphological differences among pathogens. The emergence of deep learning-driven computer vision provides a means to overcome this limitation by leveraging rich textural and morphological information directly from microscopy images^14,15^. Several strategies that combine microscopy with deep learning have been explored for pathogen identification. One common strategy is to employ standard clinical staining methods (e.g., Gram or Ziehl-Neelsen staining) to enhance contrast and introduce taxonomically informative color features^16,17^; When combined with deep learning, stain-enhanced approaches have shown promising results and, in some cases, have progressed to clinical testing^18^; the added color dimension, however, is both its strength and an operational constraint, as it necessitates additional reagents, processing time, and standardization across laboratories. An alternative route replaces chemical dyes with fluorescent probes targeting specific molecular signatures, exemplified by fluorescence in situ hybridization (FISH) against rRNA sequences. This achieves high taxonomic specificity through direct molecular recognition^19^, but at the cost of requiring fluorescence microscopy and facing inherent limitations in multiplexing capacity. Raman spectroscopy circumvents exogenous labels altogether by harnessing intrinsic vibrational signatures. When paired with deep learning, Raman-based imaging enables label-free, information-dense identification at the single-cell level^20^. Despite this technical appeal, the dependence on specialized, costly hardware has so far limited its adoption in routine clinical workflows.

Label-free bright-field imaging with standard clinical microscopes offers an appealing resolution to these constraints, as it preserves native cellular architecture and leverages existing clinical infrastructure. However, translating this potential into a clinically accurate system confronts a critical resource gap: the development of high-performing computer vision models for object classification typically requires training datasets comprising hundreds of thousands to millions of annotated, semantically diverse target images^21,22^, yet no such large-scale dataset exists for label-free bright-field microscopy of clinically relevant bacterial pathogens. Consequently, the diagnostic capacity of deep learning in this modality remains largely uncharacterized.

Here, to overcome this data bottleneck and establish proof-of-principle for rapid, label-free ESKAPe (e for *E coli* substituting *Enterobacter spp.*) identification using standard optical microscopy, we developed a cost-effective, high-throughput bacterial imaging platform and acquired over 24 million single-bacterium snapshots. Using this resource, we trained a deep neural network to classify ESKAPe pathogens based on their microscopic morphological features at the single-cell level. We further explored feasibility in two contexts: with reference strains in simulated clinical matrices, and with native clinical specimens (respiratory specimens and blood) after brief cultivation to enrich bacterial biomass. In both cases, we achieved accuracy comparable to sequencing-based methods while substantially reducing turnaround time relative to standard culture-based identification workflows. While not yet a culture-free diagnostic, this workflow establishes the proof-of-concept and foundational resources necessary for future culture-free deployment, and presents a comprehensive, end-to-end pipeline spanning data collection, model training, and preliminary clinical exploration (Figure 1).

**Figure 1.**
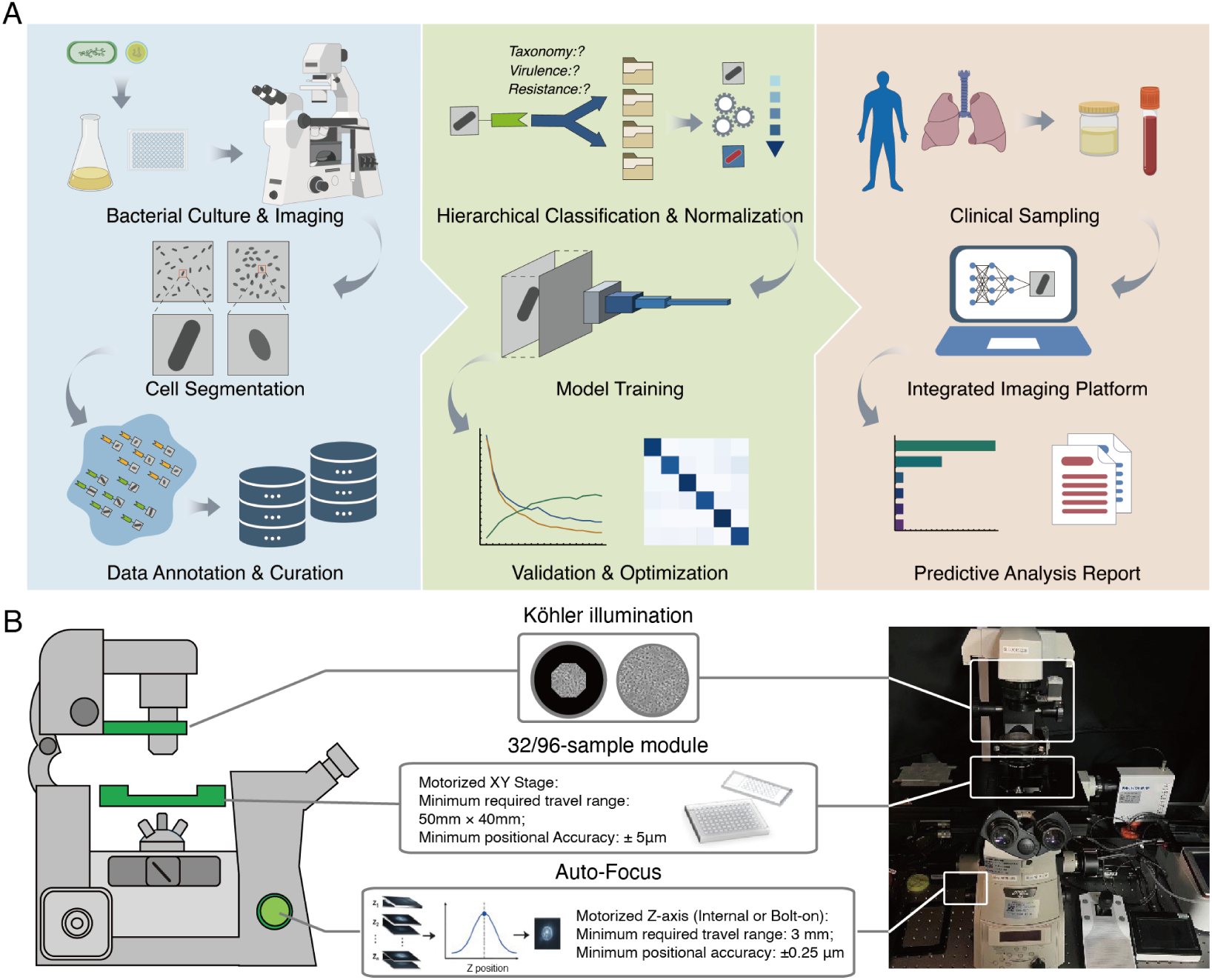
Workflow of the bacterial identification based on optical imaging and deep learning. Our system is composed of three sequential modules. (A) **Data acquisition.** Using optical microscope and custom-made gadgets, ESKAPe pathogens and other bacterial species are cultured under various conditions and incubation times, followed by optical microscopy imaging and automatic data acquisition. Subsequent image processing involves cell segmentation, manual annotation, and curated dataset assembly for deep learning applications. **Training of image deep-learning model**. The raw data are classified and normalized, followed by ResNet50 convolutional neural network training with hyperparameter optimization to optimze classification accuracy. **Clinical application.** Specimens (sputum and blood samples) are collected and processed with the same imaging platform, and then analyzed by the deep-learning model for bacterial identification. (B) Automated imaging setup used for dataset acquisition. Imaging was performed on an inverted microscope equipped with a Dhyana 401D sCMOS camera (6.5 μm pixel size; 2048 × 2048 pixels; 16-bit) and a 60× oil-immersion objective (NA 1.42), with an additional 1.5× intermediate magnification (total magnification, 90×; effective pixel size, 72.2 nm). Köhler illumination was adjusted prior to acquisition to ensure uniform illumination and optimal single-cell contrast. High-throughput imaging was enabled by a motorized stage in combination with custom-designed 32- and 96-well sample holders. Autofocus was implemented using the DAPI channel to maintain focal consistency across fields of view. Photograph of the microscope system (right) used for image acquisition, showing the physical instrument configuration corresponding to the schematic model presented on the left.

## Results

### High-throughput Microscopy Imaging Set-up

To generate a sufficiently large training dataset for neural network-based classification, we constructed a high-throughput bright-field microscopy platform building upon our previously established multi-fluorescence Bacterial Cytological Profiling (BCP) system^23^. Unlike existing datasets that predominantly rely on dye-enhanced contrast (e.g., Gram or Ziehl-Neelsen staining)^24^, we focused on label-free imaging of chemically fixed pathogens. The system was built on a Nikon Ti-Eclipse inverted microscope equipped with a commercially available motorized stage. We configured the optical path using basic bright-field components (Köhler illumination, standard objectives) common to both research-grade and clinical microscopes, ensuring that imaging conditions would be translatable to widely available clinical equipment.

Key improvements focused on accessibility: we engineered a fully 3D-printable device for casting agarose pads, compatible with standard 75 × 25 mm slides and 24 × 50 mm coverslips (**Figure 1B**). This design eliminated the need for custom glass components, and the compact 32-well format reduced the required stage travel range and positional precision compared to standard multi-well plates. The resulting 32-well agarose pad is compatible with both upright and inverted microscopes equipped with Köhler illumination (**Figure 1B**).

To automate data acquisition and minimize operator-dependent variability, we developed a browser-based control software that coordinates stage motors, shutters, and cameras. This software natively implements image-based autofocus and multi-well scanning (**Figure 1B**)^25,26^. Upon validation on the Nikon Ti-Eclipse inverted microscope, the system achieved a throughput of approximately 50 seconds per well, capturing 25 images per well. To ensure biosafety compliance and standardize imaging conditions, all pathogen samples were chemically fixed with equal volume 4% paraformaldehyde (PFA), following a protocol adapted from a recent report that demonstrated paraformaldehyde outperformed alternative fixatives in preserving bacterial morphology for image-based analysis.

### Microscopy Dataset Construction for ESKAPe Pathogens

Having established this high-throughput platform, we proceeded to construct a large-scale, comprehensive image dataset of six pathogens of primary clinical concern in sepsis and severe respiratory infections. Our panel comprises five ESKAPE species—*Enterococcus faecium* (*Efm*), *Staphylococcus aureus* (*Sau*), *Klebsiella pneumoniae* (*Kpn*), *Acinetobacter baumannii* (*Aba*), *Pseudomonas aeruginosa* (*Pae*) —with *Escherichia coli* (*Eco*) substituted for *Enterobacter spp.* due to its higher isolation frequency in the studied clinical setting, and a lack of standard species/strains for *Enterobacter spp.*^27^ (**Figure 2A**). To ensure the downstream neural network models learned robust, generalized species features rather than strain-specific or growth-phase-dependent artifacts, we incorporated intra-species biological variation (multiple strains, growth phases) and technical variation (imaging sessions, slide preparations). Our dataset encompasses multiple distinct strains per species: eight strains for *Kpn*, two for *Aba*, three for *Pae*, and one each for *Efm*, *Sau*, and *Eco*. To capture dynamic morphological changes across bacterial life cycles, each strain was cultured in Luria-Bertani (LB) broth and harvested at three distinct growth stages: early exponential (OD600 ≈ 0.1), mid-exponential (OD600 ≈ 0.5), and stationary (OD600 ≈ 1.0).

**Figure 2.**
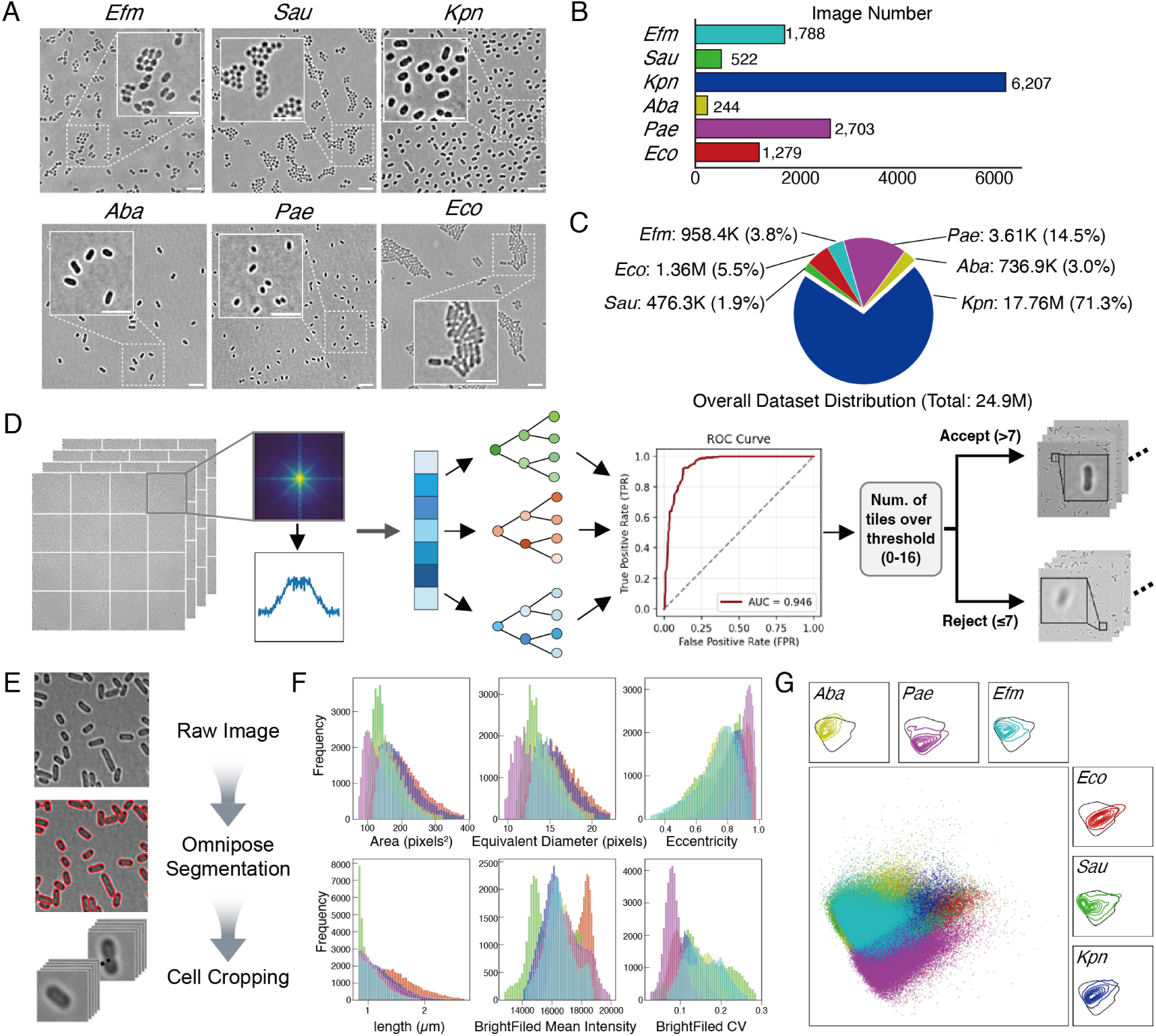
Dataset composition, image quality control, and single-cell feature profiling of the ESKAPe microscopy dataset. (A) Representative bright-field images illustrating the morphological characteristics of bacteria in the ESKAPe dataset, including *Enterococcus faecium* (*Efm*), *Staphylococcus aureus* (*Sau*), *Klebsiella pneumoniae* (*Kpn*), *Acinetobacter baumannii* (*Aba*), *Pseudomonas aeruginosa* (*Pae*) and *Escherichia coli* (*Eco*). (B) Distribution of image counts across species, showing the overall scale and class composition of the dataset at the image level. (C) Distribution of total single-cell counts for each bacterial species, reflecting dataset scale at single-cell resolution. (D) Automated image quality control pipeline. Each acquired image is partitioned into a 4 × 4 grid of panels, followed by feature extraction in the frequency domain via Fourier transformation. A random forest classifier is then applied to assign a focus-quality score to each panel. Panels exceeding a predefined threshold are retained, whereas low-scoring panels are excluded, thereby removing out-of-focus regions and improving dataset quality. Classifier performance is evaluated using a receiver operating characteristic (ROC) curve. (E) Single-cell segmentation and extraction. Raw microscopy images are processed using Omnipose to generate segmentation masks, followed by cropping of individual cells for downstream quantitative analysis. (F) Distributions of representative single-cell features across bacterial species, including morphological (e.g., area, equivalent diameter, eccentricity, and cell length) and intensity-based parameters, highlighting species-specific phenotypic variability. (G) Principal component analysis (PCA) of single-cell morphological feature profiles across the ESKAPe dataset. Each point represents an individual bacterial cell projected onto the first two principal components based on morphological and intensity-derived features. PCA shows limited linear separability among species. Kernel density estimation (KDE) contours also show no distinguishable differences in distribution patterns within the embedding space.

Raw bright-field micrographs were processed through an automated computational pipeline to extract high-quality single-cell crops for deep learning training. First, to filter out out-of-focus artifacts inherent to high-throughput slide scanning, we implemented an automated focus quality evaluation: raw images were divided into a 4 × 4 grid of 16 patches, and regional frequency profiles were extracted via Fast Fourier Transform (FFT). A random forest classifier, trained on manually annotated patches, achieved robust performance (ROC-AUC = 0.946) in discriminating in-focus regions (**Fig. 2D; see Methods**). Images with at least seven in-focus patches—empirically determined to balance yield and quality—were accepted for downstream analysis. Approved images were then processed for cellular segmentation using Omnipose^28^ and custom post-processing scripts, generating precise semantic masks and bounding boxes for cropping individual bacteria (**Figure 2E**). This pipeline yielded a large, curated dataset of 24.9 million single-cell image crops derived from over 10,000 raw fields of view (**Figure 2B, C**). Preliminary analysis of this single-cell dataset revealed extensive overlap in primary morphological features—such as cell area, equivalent diameter, and length—across the studied species (**Figure 2F**), which highlights the necessity of neural network-based approaches for reliable classification.

### Neural Network Training and Validation on Pure Cultures

We used neural networks developed for image recognition to establish a proof-of-concept identification model for our target pathogen panel, based on cell images from pure cultures. After intensity normalization and mirror padding to 224 × 224 pixels to conform to ResNet50 input specifications (Methods), we employed ResNet50 (Residual Network)^29^ as the backbone architecture and trained a model using 80% of the bacterial cell images, with 10% reserved for testing and 10% for validation. During model training, we performed hyperparameter optimization, adjusting learning rate, and batch sizes; early stopping based on validation loss was employed to avoid over-fitting. After 12 epochs, model performance plateaued on the validation set (**Figure S1**), achieving an overall accuracy of 0.92 (F1 score and ROC-AUC were 0.92 and 0.99, respectively) for species-level classification (**Figure 3A**). This model is hereafter referred to as ESKAPe-ResNet. The confusion matrix revealed that the model achieved the highest per-species recall for *Aba* (97.9%) and the lowest for *Sau* (87.3%). Among misclassified Sau cells, 4.35%, 4.31%, 1.85%, 0.97%, and 1.22% were misidentified as *Efm*, *Kpn*, *Aba*, *Eco*, and *Pae*, respectively (**Figure 3B**). A normalized confusion matrix on the held-out test set is provided in **Figure S2**. We further evaluated the reliability of ESKAPe-ResNet by analyzing prediction confidence and uncertainty. Correct predictions exhibited substantially higher maximum predicted probabilities and lower entropy than incorrect predictions, with distinct species-specific confidence profiles (**Figure S3**).

**Figure 3.**
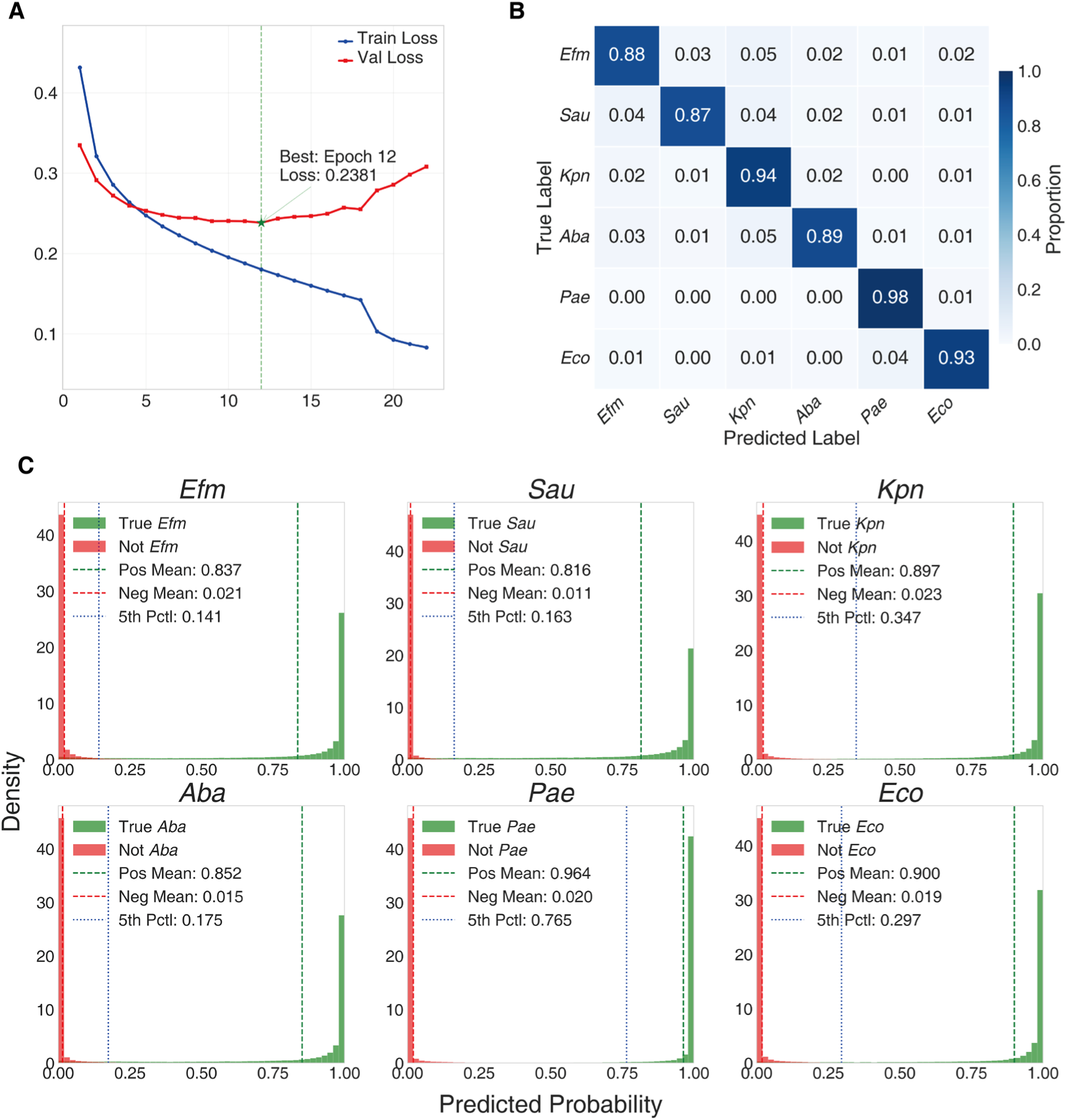
Training dynamics and performance evaluation of the ESKAPe-ResNet model. (A). Training and validation loss curves across epochs. The optimal model was achieved at epoch 12 with a minimum validation loss of 0.2381 (green dashed line). (B). Normalized confusion matrix of the optimal model on the validation set. (C). Predicted probability distributions for the six ESKAPe pathogen classes (*Efm*, *Sau*, *Kpn*, *Aba*, *Pae*, *Eco*). Density distributions are shown for true positive instances (green) versus negative instances (red). Green and red dashed lines indicate mean probabilities of positive and negative instances, respectively; blue dotted lines mark the 5th percentile of positive distributions.

We employed t-SNE (t-distributed Stochastic Neighbor Embedding) to visualize the clustering structure of high-dimensional feature representations extracted by the network and to query how species-specific morphological information is encoded in the model’s latent space. We extracted 2048-dimensional feature vectors from the pre-classification layer of the trained ESKAPe-ResNet model. Following stratified random sampling on a large-scale cell image dataset containing six species, we first reduced the dimensionality to 50 principal components via Principal Component Analysis (PCA), then mapped the features to two-dimensions using t-SNE. The results indicated that single cells are faithfully clustered by their species of origin. Notably, this clustering does not correspond to Gram staining classification (G+ vs G-), suggesting that the model discriminates based on species-specific morphological nuances beyond cell wall architecture.

In terms of intra-species heterogeneity, we quantified intra-cluster dispersion using mean pairwise Euclidean distance, and assessed clustering quality using silhouette scores in the t-SNE embedding space. *Efm* exhibited the highest morphological heterogeneity (69.12) and poorest clustering quality (−0.28), whereas *Kpn* formed the most compact cluster (32.85) with the highest silhouette score (0.40) despite inclusion of the most strains. Overall, the results demonstrate that ESKAPe-ResNet reliably distinguishes between species and capture their morphological variations.

To examine the selectivity of ESKAPe-ResNet, we evaluated five non-ESKAPe, common commensal gut bacterial strains from human and mice: *Akkermansia muciniphila* (*Akk*), *Bifidobacterium pseudolongum* (*Bif*), *Enterococcus faecium* ATCC 6057 (*Efm*(ATCC 6057); a probiotic strain), *Lactobacillus acidophilus* (*Lac*), and *Lactobacillus reuteri* (*Lre*) as out-group controls. We evaluated the ESKAPe-ResNet model on these commensal bacteria under forced closed-set classification (i.e., requiring assignment to one of the six training categories) to assess baseline misclassification patterns. Despite this constraint, analysis of softmax probability distributions revealed that non-ESKAPe strains received low prediction probabilities across all ESKAPe categories (maximum 0.295), indicating that the model exhibited characteristic uncertainty rather than overconfident false assignment (**Figure 4C**). t-SNE visualization corroborated this discriminative capacity, with ESKAPe pathogens and non-ESKAPe species forming well-separated clusters in the embedding space (**Figure 4D**). Comprehensive out-of-distribution diagnostics, including three-group probability comparison and threshold-dependent rejection curves, are provided in **Figure S4**. Misclassification patterns and per-species confidence profiles for individual non-ESKAPe strains are detailed in **Figure S5**. In summary, the ESKAPe-ResNet model exhibits robust out-of-distribution detection and effectively avoids overconfident misclassification of non-ESKAPe commensals.

**Figure 4.**
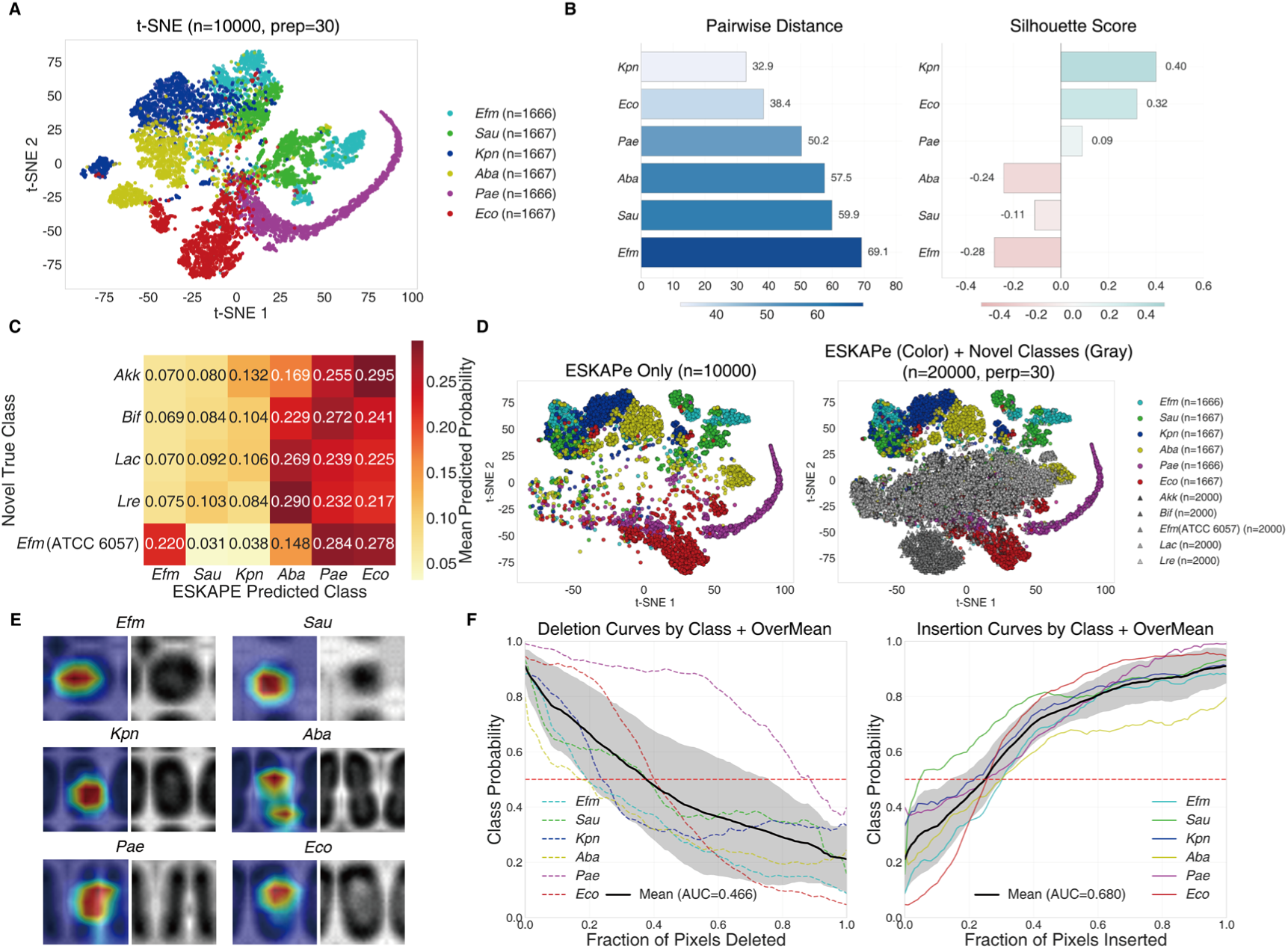
Model interpretability and generalization analysis of ESKAPe-ResNet. (A). t-SNE visualization of 10,000 ESKAPe pathogen embeddings (perplexity = 30), colored by species. (B). Quantitative assessment of bacterial morphological heterogeneities: left panel shows mean pairwise distances indicating intra-species dispersion (light to dark blue); right panel shows silhouette scores measuring cluster separability, where positive values (teal) indicate well-separated clusters and negative values (pink) suggest overlap with neighboring clusters (gray line at zero). (C). Mean predicted probability matrix for non-ESKAPe bacterial species (*Akk*, *Bif*, *Efm*(ATCC6057), *Lac*, *Lre*) forcibly classified into ESKAPe categories; warmer hues (yellow to red) indicate higher misclassification probabilities. (D) Comparative t-SNE visualization of ESKAPe pathogens (colored circles) versus non-ESKAPe species (gray triangles) at 20,000 total samples (10,000 per group); left panel shows ESKAPe-only distribution, right panel shows combined embedding space. (E). Representative Grad-CAM visualizations for the six ESKAPe classes, with red indicating high activation concentrated at bacterial centers. (F). Insertion-deletion faithfulness curves: deletion curves (left) show probability decay as important pixels are masked (lower AUC = better localization); insertion curves (right) show probability recovery as salient features are restored (higher AUC = better feature capture). Black solid lines represent mean ± standard deviation across classes; colored dashed lines indicate individual class trajectories; horizontal red dashed lines mark random chance (probability = 0.5).

**Figure 5.**
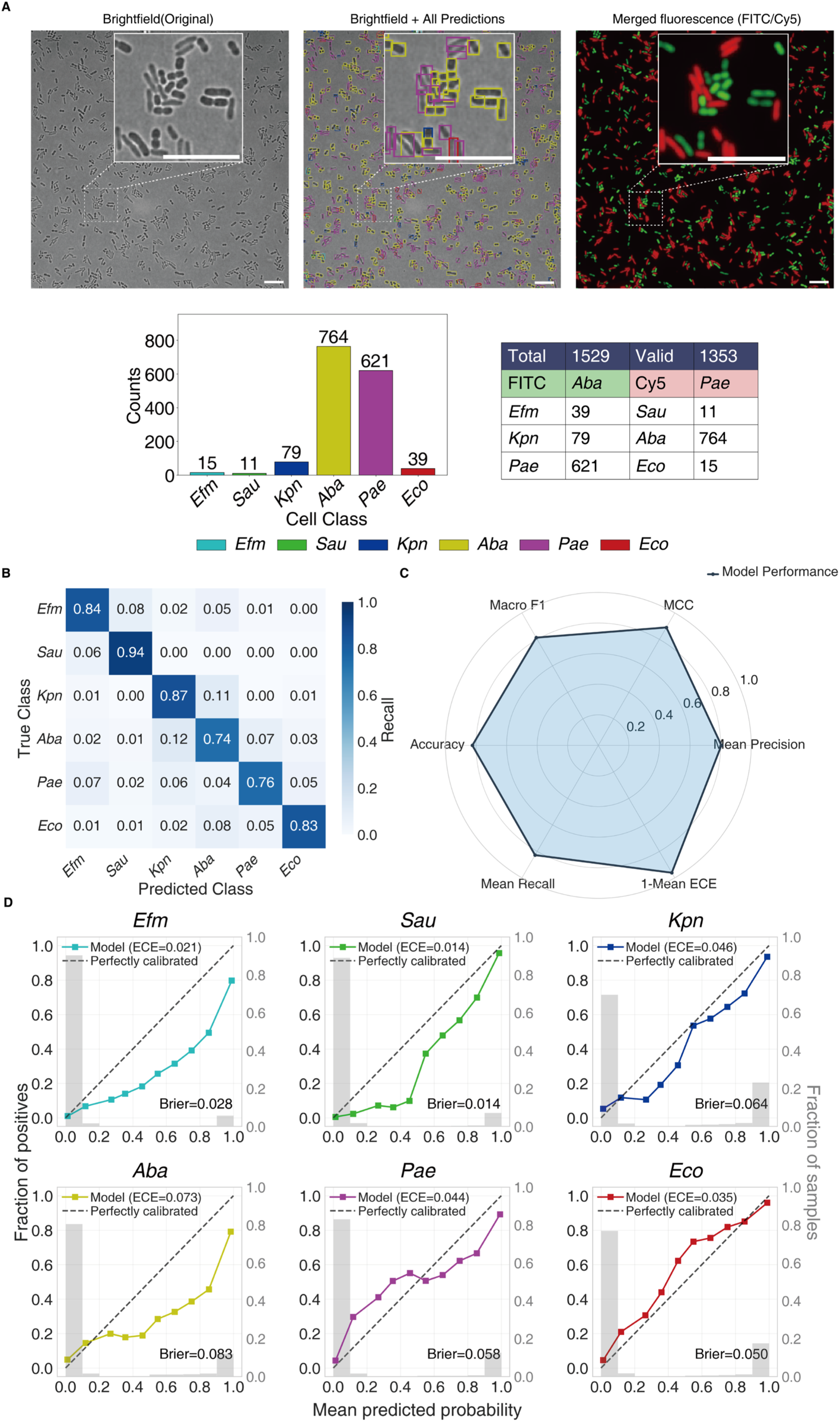
Validation of ESKAPe-ResNet on dual-fluorescence labeled bacterial mixtures. (A). Representative prediction and fluorescence validation of ESKAPe-ResNet on dual-fluorescence labeled bacterial mixtures. All panels include a magnified inset and a 100 μm scale bar (white bar, lower right). Left: original brightfield image. Middle: brightfield image overlaid with all predicted cell bounding boxes colored by predicted class (color key at bottom). Right: merged fluorescence image of FITC (green, *Aba*) and Cy5 (red, *Pae*) channels. Bottom left: bar chart showing predicted cell count distribution across six ESKAPe pathogens. Bottom right: summary statistics table showing total and valid fluorescence-labeled cell counts, the identities of FITC- and Cy5-labeled species, and predicted cell count distribution across six ESKAPe pathogens. (B). Normalized confusion matrix of cell-level predictions against fluorescence-based ground truth. (C). Radar chart displaying six comprehensive performance metrics: Accuracy, MCC (Matthews Correlation Coefficient), Macro F1, Macro Precision, Macro Recall, and 1-Mean ECE (Expected Calibration Error). (D). Reliability diagrams showing probability calibration for each ESKAPe class. Each panel displays the relationship between mean predicted probability and observed frequency of positive cases (solid line with square markers) against perfect calibration (dashed diagonal). Gray histograms indicate sample distribution across probability bins.

### Explainability of ESKAPe-ResNet assessed by Grad-CAM

To elucidate how image features extracted by ESKAPe-ResNet distinguish bacterial species, we visualized and interpreted ESKAPe-ResNet using Grad-CAM^30^; Grad-CAM uses the gradients of the target class score with respect to the final convolutional layer feature maps to quantify spatial importance of feature channels. These gradients are globally average-pooled to obtain channel-wise importance weights, which weight the forward activation maps to produce a coarse localization map. We applied Grad-CAM to the final convolutional layer of the ResNet-50 backbone (the 1×1 convolution within the last bottleneck block of stage 4, preceding global average pooling), producing 7×7×2,048 feature maps. We randomly selected 10,000 cell images (approximately 1,667 per species), computed Grad-CAM saliency maps for each image, and averaged these to obtain a species-specific activation pattern (**Figure 4E**). The resulting visualizations demonstrated that the model focuses on cell bodies rather than background regions, indicating that its predictions derived from biologically meaningful morphological features.

To quantitatively assess the faithfulness of these Grad-CAM visualizations, we conducted an insertion-deletion analysis. During the deletion evaluation, pixels were ranked by their Grad-CAM importance scores and progressively replaced with spatially Gaussian-blurred counterparts (σ = 10 pixels) to monitor the decay in classification probability (**Figure 4F**). Conversely, during the insertion evaluation, these important pixels were progressively unmasked from a blurred baseline to monitor the recovery of the model’s confidence. The near-monotonic decay observed in the deletion curves (**Figure 4F, left panel**), coupled with the progressive recovery in insertion curves (**Figure 4F, right panel**), demonstrates that the Grad-CAM attributions align with the model behavior and corroborates the faithfulness of these explanations. Per-class faithfulness metrics, including species-specific deletion and insertion curves with corresponding AUC comparisons, are provided in **Figure S6**.

### Quantifying ESKAPe Pathogens in Mock Mixed Communities

Polymicrobial infections are common in clinical specimens, and determining the relative abundance of constituent pathogens is essential for guiding targeted antimicrobial therapy. This requires identifying the species of individual cells in mixed samples and then quantifying the relative proportion of different species. Hence, we applied ESKAPe-ResNet to mock mixtures. Two species from our target panel were seperately labeled with amine-reactive fluorescent dyes (FITC-NHS and Cy5-NHS), then mixed at defined ratios (see Methods). We constructed 47 mock mixtures encompassing pairwise combinations of the six species at varying ratios, and acquired both bright field and dual-fluorescence microscopy images. In total, we acquired 460 fields of view, yielding approximately 460,000 cells for analysis downstream analyses.

With fluorescent labels serving as ground truth for individual cell identity, we evaluated ESKAPe-ResNet’s single-cell classification performance in mock mixtures. For each mixture, we compared model predictions against fluorescent labels on a cell-by-cell basis. The model achieved a pooled classification accuracy of 82.41%, macro-averaged F1 score of 0.8137, and Matthews Correlation Coefficient (MCC) of 0.7808 across all 47 mock mixtures. Per-class precision, recall, and F1-score, together with the relationship between calibration error and classification accuracy for each species, are detailed in **Figure S7**.

Expected Calibration Error (ECE) measures the agreement between predicted confidence and actual accuracy and we calculated per-species ECE. The lowest ECE was for *Sau* at 1.38%, and for *Efm* it was 2.07%, indicating highly consistent predicted probabilities with actual abundances; for *Aba* ECE was 7.29%, indicating mild overconfidence. Another measure, Brier score is the mean squared error of probabilistic predictions, and for the six species we calculated an average of 0.049, reflecting the overall accuracy of the model’s probability outputs. These metrics indicate that in the mock communities, ESKAPe-ResNet achieves accurate and statistically reliable species identification at the single-cell level.

### Proof-of-Principle Validation on Positive Blood Cultures

To evaluate the value of our workflow for future clinical deployment, we applied label-free optical imaging and ESKAPe-ResNet to positive blood culture broths from sepsis patients, and assessed the model’s ability to identify target pathogens. To this end, we collected 8 positive blood culture broths flagged by the BacT/ALERT Virtuo system (bioMérieux, Marcy-l’Étoile, France, based on CO₂ production detected by colorimetric sensors)^31^, then simultaneous acquired microscopy images using the aforementioned imaging platform and conducted 16S rRNA amplicon sequencing. Using 16S rRNA sequencing results of the same culture broths as references, we found seven samples were dominated by *A. baumannii* (highest relative read abundance), in which ESKAPe-ResNet correctly identified *Aba* as the predominant species in five samples, with 52.0%-97.6% of predicted cells classified as *Aba* (**Figure 6A, Figure S8**). ESKAPe-ResNet also correctly identified the rest one sample to be dominated by Pae **(Figure S9)**, thus achieved overall 75% accuracy. Detailed brightfield microscopy images, species-level prediction visualizations, and statistical summaries for individual blood culture samples are provided in **Figures S8–S14**.

**Figure 6.**
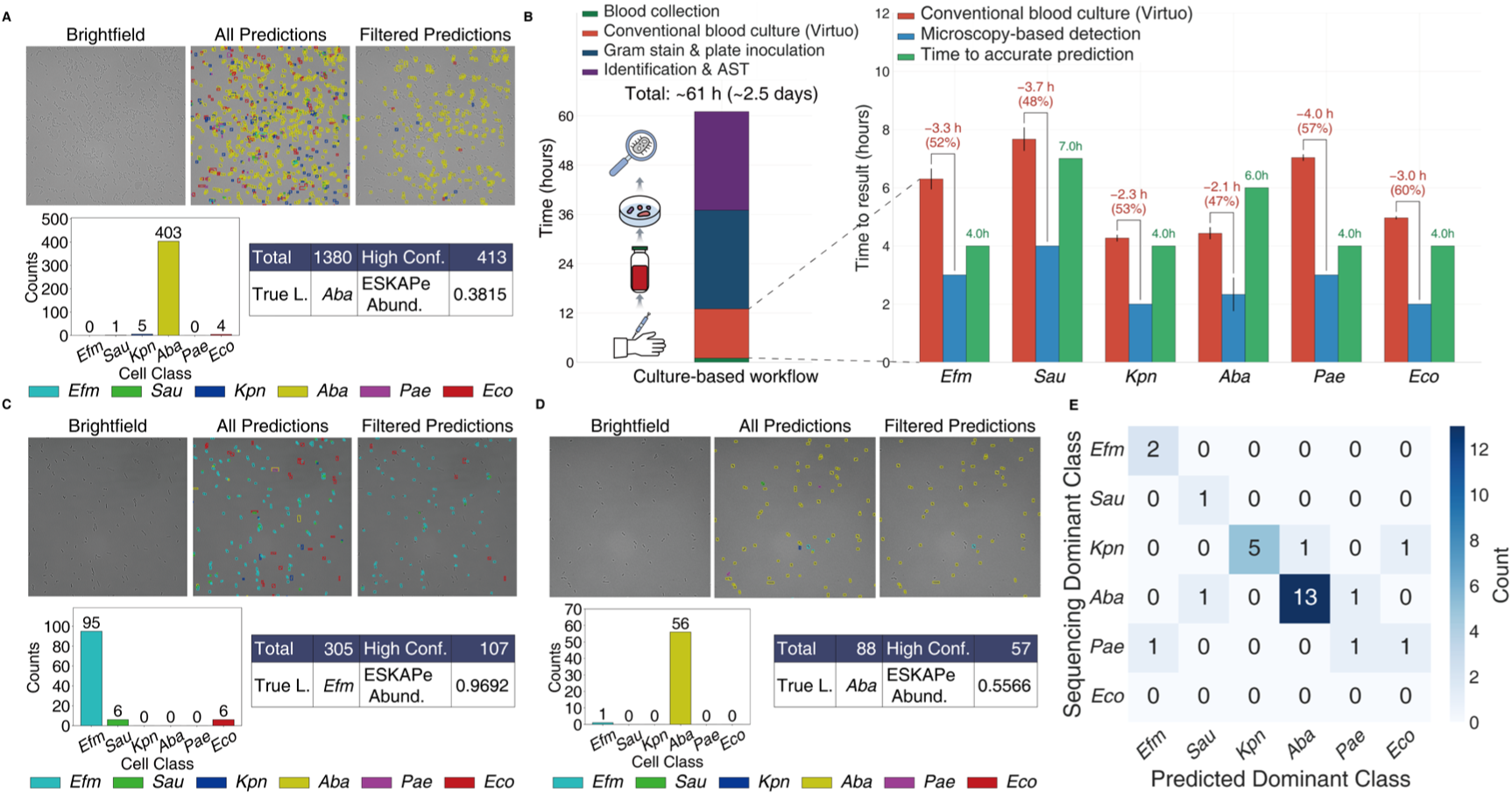
Detection of ESKAPe pathogens from clinical samples. (A) Brightfield microscopy and model prediction analysis of a blood culture sample. The top row shows the original brightfield image (left), predictions for all detected cells with bounding boxes color-coded by species (middle), and high-confidence predictions at a probability threshold of 0.998 (right). The bottom row shows the species distribution of high-confidence cells as a bar plot (left) and a statistical summary table (right) reporting the species distribution of high-confidence predictions and the ESKAPe abundance from sequencing. (B) Left, schematic of the conventional blood culture workflow showing four sequential stages: blood collection, automated blood culture incubation (with BioMérieux Virtuo), Gram staining and plate inoculation, and identification with antimicrobial susceptibility testing (AST), total time-to-result is indicated above the bar. Right, time-to-result comparison of conventional blood culture (BioMérieux Virtuo, red bars), microscopy-based detection (blue bars), and time to accurate prediction (green bars) for six ESKAPe species. Data are means ± standard deviation from three independent replicates. Brackets indicate absolute and relative time reduction of microscopy-based detection relative to conventional blood culture. Values above green bars represent the mean time to achieve accurate prediction, defined as the earliest time point with >70% classification accuracy. (C) Sputum sample analysis in the same layout as (A). (D) Bronchoalveolar lavage fluid (BALF) sample analysis in the same layout as (A). (E) Confusion matrix of dominant pathogen classification across 28 clinical samples (blood, sputum, and bronchoalveolar lavage fluid), comparing model predictions with sequencing results. Rows correspond to the ground-truth dominant class determined by sequencing, and columns correspond to the model-predicted dominant class. Numbers in each cell represent sample counts.

To further investigate the potential of our pipeline to shorten time-to-identification, we carried out a time-course experiment using spiked blood culture broths inoculated with reference strains. One representative strain from each target species was inoculated separately into standard clinical blood culture media. Cultures were sampled hourly for analysis using our approach. Parallel replicates using the same strains and media were monitored with BacT/ALERT (**Figure 6B**). BacT/ALERT flagged positive growth after 4.2–7.9 hours (mean 5.8 hours), based on CO₂ accumulation detected by colorimetric sensors. Applying ESKAPe-ResNet, we correctly identified pathogens inoculated in these blood cultures (85.19% of the cells were correctly classified to the inoculated ESKAPe species), despite the drastic change in growth media. Moreover, our approach reduced turnaround time for species-level identification by 2.1–4.0 hours (47%–60% reduction; mean 3.1 hours) relative to BacT/ALERT positivity alone—though additional time for downstream species identification (e.g., MALDI-TOF MS) would further widen this gap in routine clinical workflows. Species-and time-resolved prediction accuracy in spiked blood cultures, evaluated across all segmented cells and high-confidence subsets, is shown in **Figure S15**. In summary, the results support the potential of our approach to accelerate species-level identification in positive blood cultures, establishing proof-of-concept for future integration into clinical microbiology workflows. Clinical metadata, sequencing results, and model predictions for blood culture samples are detailed in **Table S1**.

### Proof-of-Principle Validation on Respiratory Specimens

Finally, we examined our workflow’s performance on respiratory specimen, namely sputum (n=17) and bronchoalveolar lavage fluids (BALF, n=3), from patients diagnosed with hospital-acquired pneumonia or community-acquired pneumonia (HAP/CAP). To obtain sufficient bacterial counts and minimize interference from host cells and mucus, from these specimens were diluted 2- to 10-fold then cultured in BHI medium for 2-6 hours to recover viable organisms. Sample densities after short-term recovery culture were estimated to be between 10^4^ – 10^5^ CFUs/mL culture media (Figure 6C, D). For comparison, 16S rRNA sequencing was performed in parallel. We compared the top-ranked species by 16S rRNA relative read abundance with the dominant species predicted by ESKAPe-ResNet. Among 20 samples, the top-ranked species agreed in 16 cases (80.0%), among them the top-ranking concordance was highest for *A. baumannii* (83.0%) and lowest for *K. pneumoniae* (59.4%). Further analysis showed that in the remaining four discordant samples, prolonged culture markedly altered the sequencing abundance profiles, with *Aeromonas* and *Bacillus* replacing the original ESKAPe pathogens as the dominant genera and likely explaining the model’s misclassification in these cases. The distribution of ESKAPe relative abundance across clinical samples, as determined by 16S rRNA sequencing, is shown in **Figure S16**. Per-class rank deviations between model-predicted and sequencing-derived abundance rankings are detailed in **Figure S17**. Detailed brightfield microscopy images, species-level prediction visualizations, and statistical summaries for individual sputum and BALF samples are provided in **Figures S18–S35**. Clinical metadata, sequencing results, and model predictions for respiratory samples are also included in Supplementary **Table S1**.

Since timely species-level pathogen identification is for guiding antimicrobial therapy, we additionally compared turnaround times for our approach, conventional culture, and 16S rRNA sequencing on respiratory specimens. From obtaining the sample for imaging using our platform to analytical results by ESKAPe-ResNet, we reached an average time of 4.3 ± 1.7 hours (median 5.0 hours; range 2–6 hours) for respiratory samples, in which from fixation of liquid culture to obtaining identification results took 30 minutes on average. In comparison sequencing took >36 hours on Illumina platform from DNA extraction, library preparation and obtaining sequencing results. The pronounced reduction in turnaround time for species-level identification in respiratory specimens suggests the potential of combining label-free imaging and deep-learning to accelerate pathogen detection workflows.

## Discussion

In this work, we conducted a proof-of-concept study demonstrating that standard bright-field microscopes, augmented with minimum accessories, can enable deep learning-assisted pathogen identification with potential for future clinical deployment. This workflow includes acquiring large-scale image datasets for bacterial pathogens (millions of cells) using affordable hardware adaptable to standard microscopes; training a species-specific neural network model operating at the single-cell level; and finally evaluating the pipeline on respiratory infection and sepsis samples, with a focus on turnaround time and species-level accuracy. This relatively straightforward workflow is readily transferable to clinically used microscopes, extensible to additional pathogenic species and strains, and potentially applicable to a broader range of clinical specimen types. In this regard, our workflow avoids the need for standardized staining, high-end fluorescence microscopes or other resource-intensive equipment (e.g., Raman-spectroscopy)^20^, instead leveraging standard light microscopes already available in clinical settings with affordable accessory hardware (e.g., camera and motorized stage); identification can be then carried out on a GPU-equipped laptop or remotely on a server.

Deep learning model performance is frequently constrained by training data scarcity—the so-called “scaling law” of machine learning^32,33^. We addressed this throughput bottleneck by integrating affordable hardware accessories with standard microscopes and custom bioinformatic pipelines, enabling rapid acquisition of millions of single-cell images. Nevertheless, we must acknowledge that our study panel departs from the canonical ESKAPe group by substituting *E. coli* for Enterobacter spp. This modification reflects the epidemiological reality of our clinical setting, where *E. coli* predominates among Enterobacterales isolates from bloodstream and respiratory infections^27^. While this limits direct comparability to studies employing the strict ESKAPe definition, it enhances the clinical relevance of our findings for the patient population under study. Future work will necessarily expand the training dataset to include not only *Enterobacter spp. —*so to validate generalizability across the full ESKAPe spectrum*—*but also other commensal and pathogenic microbes of interest^34–36^. There in constructing a foundational vision model for label-free bacterial identification, the core challenge lies not in model architecture but in systematic, large-scale data curation.

Our study has demonstrated the proof-of-principle that deep visual models (e.g., ResNet) can achieve robust pathogen identification from standard bright-field micrographs. As visual model architectures and training strategies continue to evolve^37–39^, several avenues emerge for optimizing this pipeline toward clinical deployment. One critical direction is the reduction of model complexity: such optimization is critical for edge computing, enabling the deployment of these models directly onto point-of-care testing (POCT) devices and standard laboratory hardware to further accelerate diagnostic turnaround^40,41^. Furthermore, as the training dataset expands to encompass a broader diversity of species, strains, and clinical culturing conditions, re-training models from scratch will become computationally prohibitive. Adopting transfer learning or continual learning strategies to fine-tune existing models will simplify the scalable expansion of our diagnostic framework^42,43^. Finally, exploring state-of-the-art computer vision architectures—such as Vision Transformers (ViTs) and their lightweight derivatives^44–46^—offers opportunities to maximize the utility of our massive, label-free dataset. The comparative performance of these emerging architectures in extracting pathogen-specific features from diffraction-limited micrographs remains an important area for future investigation.

The inherent complexity of clinical specimens from infectious diseases necessitates specialized sample preparation and data analysis protocols, which is a critical focus for future optimization of our pipeline. One major challenge is interference from host cells, particularly in blood and respiratory specimens (sputum and BALF). Bacterial enrichment protocols—such as differential centrifugation, filtration, or targeted affinity capture—would likely improve image quality^47^, identification accuracy, and potentially eliminate the brief cultivation step (2-6 hours) still required in our current workflow^48^. Another technically demanding yet transformative direction is fine-tuning pre-trained models directly with uncultured, real-world clinical specimen, with single-cells annotated *in situ* via orthogonal methods such as FISH or Raman-spectroscopy, and sample-level corroboration by sequencing or subsequent culture-based identification^20,49–52^. Finally, incorporating antibiotic exposure during brief cultivation may enable simultaneous species identification and resistance phenotyping^53,54^; preliminary experiments suggest that resistance-associated morphological signatures are subtle and require dedicated, phenotypically characterized strain panels for robust prediction. Resolving these challenges is a critical next step toward clinical use.

In conclusion, we provide an easy-to-adapt workflow leveraging standard light microscopes and deep learning to achieve accurate, rapid, label-free pathogen identification in clinical specimens. By reducing time and equipment barriers, this workflow holds high potential for expansion to additional pathogen panels and broader applications in clinical and environmental microbiology.

Code availability

The python script for training the ESKAPe-ResNet and the model used in this paper, together with example images are freely available at https://github.com/IsaacYing/ESKAPe-ResNet

## MATERIALS AND METHODS

### High-Throughput Automated Fluorescence Imaging Platform

To enable large-scale bacterial cytological profiling, we developed a high-throughput automated imaging platform integrating custom-designed multi-well sample holders with a motorized fluorescence microscope, to achieve stable focal plane maintenance during sequential acquisition while minimizing cross-contamination between samples. This setup enables reproducible extraction of single-cell morphological and fluorescence features from approximately 1,000–10,000 bacterial cells per sample, with a throughput of up to 32 samples per hour.

The platform was built on a motorized Nikon Ti-E inverted fluorescence microscope equipped with a Dhyana 401D sCMOS camera (2048 × 2048 pixels, 16-bit; 6.5 μm pixel size) and a 60× oil-immersion objective (NA 1.42), together with a 1.5× intermediate magnification, resulting in a total magnification of 90× and an effective pixel size of 72.2 nm. Prior to imaging, Köhler illumination was optimized to ensure uniform illumination and high single-cell contrast. Automated image acquisition was controlled via a custom graphical user interface integrating stage movement, multichannel fluorescence imaging, and autofocus. Autofocus was performed using a two-step axial scanning strategy, consisting of a coarse scan over a 15 μm z-range followed by a fine scan over a 3 μm z-range. At each z-plane, image sharpness was quantified using the intensity coefficient of variation (CV), and the focal plane was selected based on the maximum CV value. Under the standard acquisition setting of five fluorescence channels and 16 fields of view (FOVs) per sample, imaging required approximately 3–3.5 min per sample, enabling scalable high-content phenotypic profiling across large bacterial collections.

### Image Quality Control

Images acquired from the high-throughput automated fluorescence imaging platform were first manually pre-screened and annotated. All images were firstly down-sampled by 2-fold, then divided into 4×4 non-overlapping patches of 256×256 pixels (16 patches per image). For each patch, two-dimensional fast Fourier transform (2D-FFT) was applied to convert the patch into the frequency domain. Statistical features were extracted by calculating the mean and standard deviation along both the X and Y axes of the magnitude spectrum, resulting in four 256-dimensional feature vectors per patch. These were concatenated to form a 1024-dimensional frequency-domain statistical feature vector characterizing energy distribution and directional texture of each patch.

A random forest classifier was employed for binary patch-level quality control classification, where positive samples correspond to high-quality clear images and negative samples correspond to defective images with blur, debris, or out-of-focus artifacts. Hyperparameter optimization was performed via 5-fold cross-validated grid search over the following parameter space: n_estimators ∈ {100, 200, 300}, max_depth ∈ {None, 10, 20, 30}, min_samples_split ∈ {2, 5, 10}, and max_features ∈ {’sqrt’, ‘log2’}. Based on the grid search results, the final model parameters were manually set to n_estimators = 100, max_depth = None, max_features = ‘sqrt’, and min_samples_split = 2.

Model training was conducted using an iterative self-training strategy to progressively expand the training set and enhance model robustness. An initial random forest classifier was trained on manually annotated positive and negative samples. This model was subsequently applied to unlabeled patches for prediction, and misclassified samples were identified, manually verified, and incorporated into the original training dataset. The model was retrained on the augmented dataset, with this procedure repeated for three times to refine the decision boundary and mitigate overfitting. This approach follows the self-training paradigm in semi-supervised learning, in which model predictions and manual verification provide iterative feedback for continuous performance improvement. The final classifier was trained on the fully augmented dataset and serialized using the Python joblib library for subsequent use.

For sample-level quality control screening of the full dataset, all candidate images were divided into 4×4 patches following the same procedure, and the 16 patches were classified individually by the trained random forest classifier for quality assessment. An image was considered to meet the quality control threshold and was retained for subsequent analysis if 9 or more of its 16 patches were classified as high-quality positive samples; otherwise, the image was filtered out. This strategy ensures that sample-level screening balances local quality assessment with overall fault tolerance, preventing effective samples from being discarded due to isolated patch misclassification.

### Image Preprocessing

Raw image data were saved as multi-channel TIFF files. The first channel contained the bright-field image, which was read using the python module *tifffile* for subsequent cell segmentation. Fluorescence channels were recorded when available for downstream analysis. Bright-field images were normalized using omnipose.utils.normalize99, rescaling pixel values to the 0–1 range to enhance cell boundary contrast.

### Bacterial Strains and Culture Conditions

The dataset used in this study comprised ESKAPe pathogens and related strains, including *Escherichia coli* MG1655, *Staphylococcus aureus* ATCC 43300, *Klebsiella pneumoniae* ATCC 43816, *Acinetobacter baumannii* ATCC 19606 and AB09, *Pseudomonas aeruginosa* PAO1, PA14, and PAK, and vancomycin-resistant *Enterococcus faecium* 21B188. A full list of strains is provided in Supplementary **Table S2**.

Model training data were constructed to encompass diverse cellular morphologies across growth phases. The six ESKAPe pathogens were cultivated in Luria-Bertani (LB), Brain Heart Infusion (BHI), and M9 minimal medium. Bacteria were grown in 96-well deep-well plates at 37°C with shaking at 180 rpm. Optical density at 600 nm (OD_600_) was monitored spectrophotometrically, and samples were harvested at early logarithmic (OD_600_ ≈ 0.2), mid-logarithmic (OD_600_ ≈ 0.5), and stationary (OD_600_ ≈ 1.0) phases, followed by fixation with an equal volume of 4% paraformaldehyde (PFA).

To evaluate model performance under simulated clinical conditions, ESKAPe pathogen suspensions at OD_600_ = 0.1 were inoculated into BacT/ALERT FA PLUS blood culture bottles (bioMérieux) at a 1:100,000 ratio and cultured at 37°C on a rocking platform shaker at 50 rpm. Samples were collected at 1-hour intervals from 1 to 6 h post-inoculation and fixed with an equal volume of 4% PFA. In parallel, identical inoculation conditions were used in the bioMérieux VIRTUO blood culture system at the clinical microbiology laboratory, and the time to positivity alarm was recorded for each bottle. This enabled direct comparison between the high-throughput automated fluorescence imaging platform and the standard clinical blood culture workflow.

To enable fluorescence-based discrimination of individual strains in mixed ESKAPe populations, the six ESKAPe pathogens were cultured in BHI medium in 96-well deep-well plates at 37°C with shaking at 180 rpm until reaching OD_600_ ≈ 0.8, followed by fixation with an equal volume of 4% PFA. Individual strains were differentially labeled after fixation using distinct fluorescent D-amino acid probes (FDAAs), including FITC- and Cy5-conjugated derivatives, to enable fluorescence-based discrimination between bacterial populations within the same imaging field. Labeling was performed for 10 min at room temperature in the dark, after which samples were centrifuged and resuspended in 1× PBSTw to OD_600_ = 1 prior to imaging.

Beyond the above culture systems, to assess the model’s ability to exclude non-target organisms, the dataset also included common gut commensal strains, including *Akkermansia muciniphila* DSM 26127, *Bifidobacterium pseudolongum* CGMCC1.2202, *Enterococcus faecium* ATCC 6057, *Lactobacillus acidophilus* CGMCC 1.1878, and *Lactobacillus reuteri* CICC 6131. A full list of strains is provided in Supplementary **Table S2**.

All commensal strains were inoculated into BHI medium and cultured in Hungate tubes at 37°C under anaerobic conditions until reaching OD_600_ ≈ 0.8, followed by fixation with an equal volume of 4% PFA. Prior to anaerobic cultivation, the medium was degassed in an anaerobic jar by vacuum evacuation using a pump, flushed with N₂/CO₂ (80:20) gas mixture, and sealed with a butyl rubber stopper and aluminum crimp cap after inoculation.

### Single-Cell Segmentation and Extraction

Cell segmentation was performed using Omnipose^28^. Briefly, the pre-trained *bact_phase_omni* model was deployed on an NVIDIA A100 (40GB) GPU with the following key settings: images were upsampled by a factor of 1.5 to preserve fine morphological details; both the omni and cluster functions were enabled to allow mask reconstruction and DBSCAN-based separation of adjacent cells; and *mask_threshold* and *flow_threshold* were set to zero to avoid over-filtering. After segmentation, we used *skimage.measure.regionprops* to extract bounding box coordinates for each cell, which were then applied to crop single-cell images from the bright-field channel.

### Image Standardization and Data Storage

Cropped single-cell images were resized to a fixed input size using a custom prepare_bacteria_image function. Images were resized with OpenCV Lanczos interpolation (cv2.INTER_LANCZOS4) to 224 pixels on the longest side, preserving the aspect ratio. Mirror-reflective padding (cv2.BORDER_REFLECT_101) produced fixed-size square images with the cell centered. Processed single-cell images were grouped by bacterial class according to Well ID mapping and serialized using pickle into batch files (batch size = 1000) to construct the standardized training dataset.

### Model Architecture and Training

A ResNet-50 initialized with ImageNet pretrained weights was used as the backbone. The final fully connected layer was replaced with a linear layer of six output units, corresponding to the six target pathogen classes.

The model takes 224 × 224 pixel single-cell bacterial images as input. Single-cell images were loaded as single-channel grayscale arrays (bright-field microscopy). Each image was min–max normalized to [0, 1], scaled to 8-bit unsigned integers, and converted to a three-channel pseudo-RGB format for compatibility with ResNet-50. For the training set, data augmentation included random horizontal flipping, random rotation (±10°), and random sharpening (sharpness factor 1.5, probability 0.5). All images were converted to tensors and normalized using the ImageNet mean and standard deviation. The validation and test sets were resized and normalized without augmentation.

Training was implemented in PyTorch on a server with NVIDIA A100 GPUs, using DataParallel for multi-GPU training. Datasets from multiple batches were concatenated and randomly split into training, validation, and test sets at a ratio of 8:1:1. The Adam optimizer was used with an initial learning rate of 10⁻⁴ and a batch size of 64. Unweighted cross-entropy loss was used with integer-encoded class labels. The learning rate was adjusted with ReduceLROnPlateau (patience of 5 epochs) based on validation loss. Training ran for a maximum of 50 epochs, with early stopping monitoring validation loss (patience of 10 epochs); the checkpoint with the lowest validation loss was retained. The random seed was set to 42 for reproducibility.

At the end of each epoch, the validation set was evaluated for accuracy, weighted F1 score, and multi-class AUC-ROC. Confusion matrices were saved as heatmaps every 5 epochs. The following outputs were recorded during each training run: model weights and a complete checkpoint (including optimizer state, learning rate scheduler state, and training history) saved at the lowest validation loss; a text log of the random seed, dataset paths, class-wise sample counts, and per-epoch metrics; and CSV files of training/validation loss, accuracy, F1, AUC, learning rate, and epoch timing.

The model was implemented in PyTorch, with torchvision for pretrained weights and image transforms. NumPy was used for array operations, scikit-learn for metrics (F1 score, AUC-ROC, confusion matrix), and PIL for image I/O. Data were loaded from pickle files. Matplotlib and seaborn were used for visualization. Package versions were recorded in the training environment.

### Model Interpretability Analysis

Model weights were loaded from the checkpoint with the lowest validation loss and set to evaluation mode. Images were drawn from the same batch directories used for training. For each pathogen class, up to 2,000 cells were selected by stratified random sampling to balance across sources. Each cell image was min–max normalized to [0, 1], scaled to 8-bit unsigned integers, converted to a three-channel pseudo-RGB format, resized to 224 × 224 pixels, and standardized using the ImageNet mean and standard deviation.

Grad-CAM was applied to the final convolutional layer of ResNet-50 (the 1×1 convolution within the last bottleneck block of stage 4, immediately preceding global average pooling). Forward and backward hooks were registered on this layer to capture output feature maps and the gradient of the ground-truth class logit, respectively. Gradients were globally average-pooled across channels to obtain channel-wise weights, which were multiplied with the feature maps and summed. The resulting map was passed through ReLU to produce the class activation map, which was then upsampled to 224 × 224 pixels via bilinear interpolation and scaled to [0, 1]. Grad-CAM was computed for all sampled cells.

Faithfulness was assessed using insertion-deletion experiments on a subset of the samples. A baseline image was generated by Gaussian blurring (σ = 10). Pixels were ranked in descending order by Grad-CAM importance scores. For the deletion curve, original pixels were sequentially replaced with the blurred baseline in that order; for the insertion curve, original pixels were sequentially restored from the blurred baseline. Both curves were computed over 100 steps, recording the target class probability at each step. The area under each curve (AUC) was calculated; lower deletion AUC and higher insertion AUC indicated more accurate localization. For computational efficiency, the first 50 samples underwent full evaluation, after which every 20th sample was evaluated, with a maximum of 100 samples per class. The analysis generated per-class and overall insertion-deletion curves, class-wise AUC comparisons, and CSV records of per-sample prediction probabilities and faithfulness metrics.

### Mock community benchmarking

ESKAPe bacterial cultures fixed with equal volume 4% PFA were labeled with FITC or Cy5, then mixed in pairwise combinations to generate dual-fluorescence samples. Images were acquired using the high-throughput automated fluorescence imaging platform described above, with raw data stored as multi-channel TIFF files: channel 1 for brightfield, channel 2 for FITC, and channel 3 for Cy5.

Cell segmentation was performed on the brightfield channel using Omnipose. For each segmented cell, mean fluorescence intensities in the FITC and Cy5 channels were compared against thresholds defined as the background mean plus three standard deviations. Cells were classified by their fluorescence pattern: dual-positive, FITC-only, Cy5-only, or dual-negative. These fluorescence-based labels provided ground truth for species identification at the single-cell level.

Model predictions were compared against fluorescence ground truth on a per-cell basis. Classification performance was evaluated using accuracy, per-class precision, recall, and F1 score, along with Matthews correlation coefficient (MCC) and normalized confusion matrices. Probability calibration was assessed using expected calibration error (ECE) with 10 equal-width bins, reliability diagrams, and Brier score (scikit-learn).

### Clinical Sample Analysis

The trained ESKAPe-ResNet model was applied to single-cell images from clinical samples. For each sample, cells were first segmented from brightfield images using Omnipose. Each segmented cell was then processed by the model to obtain six-class probabilities. Predictions with confidence scores below 0.998 were filtered out. The final sample-level prediction was determined by majority vote of predicted classes among high-confidence cells, with class counts and relative abundances recorded for downstream comparison.

To assess agreement between model predictions and molecular sequencing results, predicted relative abundances were compared with paired sequencing data. Concordance was evaluated using Kendall’s Tau and Spearman’s Rho for rank correlation across the six classes. Top-1, Top-2, and Top-3 consistency rates and Jaccard similarity were calculated to evaluate agreement in identifying the dominant pathogen. Exact rank match distributions and per-class rank deviations were analyzed to assess the model’s ability to reproduce the relative ordering of all six classes. Additionally, the total relative abundance of ESKAPe pathogens in each sequencing sample was computed to examine the effect of low-abundance samples on model concordance.

### Clinical Sample Collection and Processing

Clinical sputum and bronchoalveolar lavage fluid (BALF) samples: Specimens were first diluted 2- to 10-fold in 1× PBS (dilution factor adjusted according to sputum viscosity). An aliquot of 5 μL of the diluted sample was inoculated into 500 µL BHI medium at a 1:100 ratio in 96-well deep-well plates and cultured at 37°C with shaking at 180 rpm. Samples were collected at 1-hour intervals from 1 to 6 h post-inoculation and fixed with an equal volume of 4% PFA.

Clinical blood samples: Blood culture bottles were vented using a 0.22 μm filter membrane, and 400 μL of blood was aspirated and mixed with 2 volumes (800 μL) of 4% PFA, followed by incubation for 15 min. To lyse erythrocytes and reduce background, Triton X-100 was then added to a final concentration of 0.2%, followed by incubation for 10 min and centrifugation at 5,000 × g for 4 min. The supernatant was discarded, and the pellet was washed twice with PBS (0.2% Triton X-100). The cell pellet was then resuspended in 1 mL of 4% PFA.

For imaging, 2% (w/v) agarose prepared in 1× PBS was melted and cast into a custom-designed imaging mold. After the agarose completely filled the wells, a clean glass coverslip was gently applied to generate a flat imaging surface while minimizing air bubble formation. Following gel solidification, stained bacterial suspensions were deposited onto the agarose pads and covered with a second coverslip for immobilization during imaging. The custom mold design enabled physical separation of samples while maintaining a uniform focal plane during automated acquisition.

All clinical samples were subjected to 16S rRNA gene sequencing using Illumina platform for species identification and relative abundance quantification.

### Software environment and dependencies

All computational analyses were performed in Python 3.8.20 with PyTorch 1.13.1 (CUDA 11.6, cuDNN 8.4.0) using NVIDIA A100 (40GB) GPUs with CUDA acceleration. Key dependencies included torchvision 0.14.1, NumPy 1.24.4, scikit-learn 1.3.2, Pillow 10.4.0, matplotlib 3.7.5, and seaborn 0.13.2. Cell segmentation was performed using Omnipose 1.0.6.

